# Spatiotemporal cellular dynamics of the notochord shape intervertebral disc morphogenesis in the mouse embryo through apoptosis and proliferation

**DOI:** 10.1101/2025.09.02.673643

**Authors:** Julie Warin, Estelle Balissat, Pauline Colombier, Lily Paillat, Maeva Dutilleul, Anne Camus

## Abstract

**Background:** The notochord is a midline structure essential for vertebrate embryogenesis, contributing to the development of the nervous system, digestive tract, and vertebral column. In particular, notochord signaling is indispensable for proper patterning and coordinated development of alternating vertebrae and intervertebral discs (IVDs). Later, notochordal cells (NCs) mature and adopt a characteristic vacuolated morphology before giving rise to the core of the forming IVD, the nucleus pulposus (NP). Postnatally, NCs play pivotal role in maintaining disc integrity through the secretion of specific factors and extracellular matrix (ECM). Despite its importance in disc formation and homeostasis, the morphogenetic mechanisms underlying the notochord’s transformation into the NP are insufficiently characterized.

**Results:** We conducted a comprehensive histological and immunohistochemical analysis to investigate the cellular events governing NP formation in the mouse developing spine. Temporal analysis of intracytoplasmic vacuole formation using *Lamp1* marker revealed their contribution to NP growth, while cell density progressively decreased. In addition, quantitative analyses demonstrated a notable proliferative capacity within notochordal cells coupled with region-specific apoptotic activity in the sclerotome, at future disc sites.

**Conclusions:** This study highlights the intricate balance of cellular proliferation, programmed cell death, matrix remodeling, and vacuolation dynamics as key determinants in shaping the NP along the rostro-caudal axis.

**Key Findings:**

- Spatiotemporal cellular changes drive the transition from notochord to nucleus pulposus
- Future disc regions show selective notochord proliferation and sclerotome cell death
- Notochord vacuolization and matrix deposition contribute to nucleus pulposus morphogenesis

## Introduction

Notochordal cells (NCs) are a rare cell type present at the midline along the anterior-posterior axis of all chordate embryos. In the early stage, NCs are organized into a mesodermal rod-like structure, the notochord, which, beyond serving a structural function, play key roles in signaling and in orchestrating the development of surrounding tissues in vertebrates. These include the central nervous system, axial skeleton, and foregut endoderm. Abnormal notochordal development is associated with a variety of birth defect in human, impacting neural, mesodermal and endoderm derivatives. A substantial number of animal studies have demonstrated the essential role of the morphogen Sonic Hedgehog (SHH), secreted by NCs, in the correct dorso-ventral patterning and differentiation of the neural tube and somites, primarily through its gradient-dependent signaling mechanism that regulates gene expression in target cells^1,2^. In somites specifically, once the ventral-medial part differentiates into sclerotome, SHH signal will promote their proliferation and migration around the notochord to form the unsegmented perinotochordal tube of loose mesenchymal sclerotome-derived cells. After their migration is completed, vertebral segmental patterning takes place through sclerotome segmentation, during which cells reorganize to form alternating condensed regions-giving rise to the annulus fibrosus (AF), the fibrous peripheral part of the intervertebral disc (IVD)-and loose regions, which develop into the vertebral bodies (VB) of the future vertebral column^2–6^. Genetic studies have demonstrated that NCs contribute to another component of the IVD, the nucleus pulposus (NP). During development NCs regress to finally reside at the center of the disc, forming the highly hydrated part of the IVD where degenerative changes are thought to initiate^7,8^. Once again, SHH signaling is critical for IVD formation, post-natal growth and for maintaining its integrity^9–12^.

Although the aetiology of low back pain is not fully elucidated, it is often associated with aging and degeneration of the disc. NC loss in certain species coincides with the onset of degenerative IVD changes. There is currently no effective treatment for disc degeneration^13^. This is largely due to a lack of basic knowledge of the molecular and cellular controls of disc differentiation, growth and homeostasis, during embryogenesis and at different stages of life. Nevertheless, there is now considerable evidence from *in vitro* studies that NCs have a significant influence on disc cells homeostasis^14–16^. At present, NCs and their biologically active factors, although promising targets for regenerative or symptom-modifying therapies for IVD disease, involve mechanisms and influencing factors that largely remain to be characterized.

Cell morphology, molecular identity and lineage-tracing experiments have established that the NP develops from the embryonic notochord. However, NCs’ cellular changes toward their functional state in the NP, defined by their intrinsic properties and specialized roles throughout developmental stages of skeletal maturation are insufficiently explored. During early organogenesis, the disc developmental process encompasses the transition from an initial state as notochord primordium (or anlage) to its mature form of the NP of the adult disc. The transition from notochord to NP involves spatiotemporal changes from a rod-like structure to segmented fibrocartilaginous tissue. Despite descriptive studies, the cellular and molecular mechanisms underlying the transformation of the embryonic notochord into the NP remain poorly understood^17,18^.

To gain insights on the morphogenetic events occurring in the notochord during IVD formation in mice, we performed histological and immunohistochemical analysis to investigate the dynamics of cellular events that orchestrate NP formation, focusing on apoptosis and proliferation. This study also examines in details cellular vacuolation and extracellular matrix (ECM) formation. These processes were analyzed along the rostro-caudal axis at key developmental stages in IVD formation, where cellular changes and complex tissue rearrangements are particularly pronounced^6,12,19–21^.

Quantitative analyses reveal a distinct capacity for cellular proliferation in the notochord, alongside evidence of differential apoptotic cell death in the sclerotome at the level of the forming disc. Our results indicate that these morphogenetic events progressively occur following a cranio-caudal pattern during axial development. Altogether our findings underscore a coordinated spatial and temporal regulation of cell proliferation and apoptosis, along with intracytoplasmic vacuoles formation and ECM remodelling as key cellular processes driving NP morphogenesis. This study provides a foundation for future investigations into the morphogenesis of the notochord and the NP in genetic mutant contexts, as well as the underlying molecular mechanisms.

## Results

### Notochord morphological changes during nucleus pulposus development

We first aimed at developing a temporal histological atlas to examine in detail the key cellular events of the notochord developing into the NP found at the core of the adult disc (Fig. 1A-O). Both Hematoxylin and eosin (H&E), and periodic acid-Schiff with alcian blue (PAS-AB) staining were performed on serial histological sections to highlight different features along developmental time from E9.5 to E17.5 and at 4 days post-natal stage (P4). H&E staining by distinguishing cell morphology and nuclei helped to reveal the general spatial organization of NCs as well as their vacuolation, and the structural changes of the surrounding tissues, in particular sclerotomes. PAS-AB staining distinguishes the ECM components and therefore was useful for highlighting the basement membrane around the notochord, as well as glycosaminoglycan (GAG)-rich regions which become more prominent as cells differentiate.

**Figure 1.**
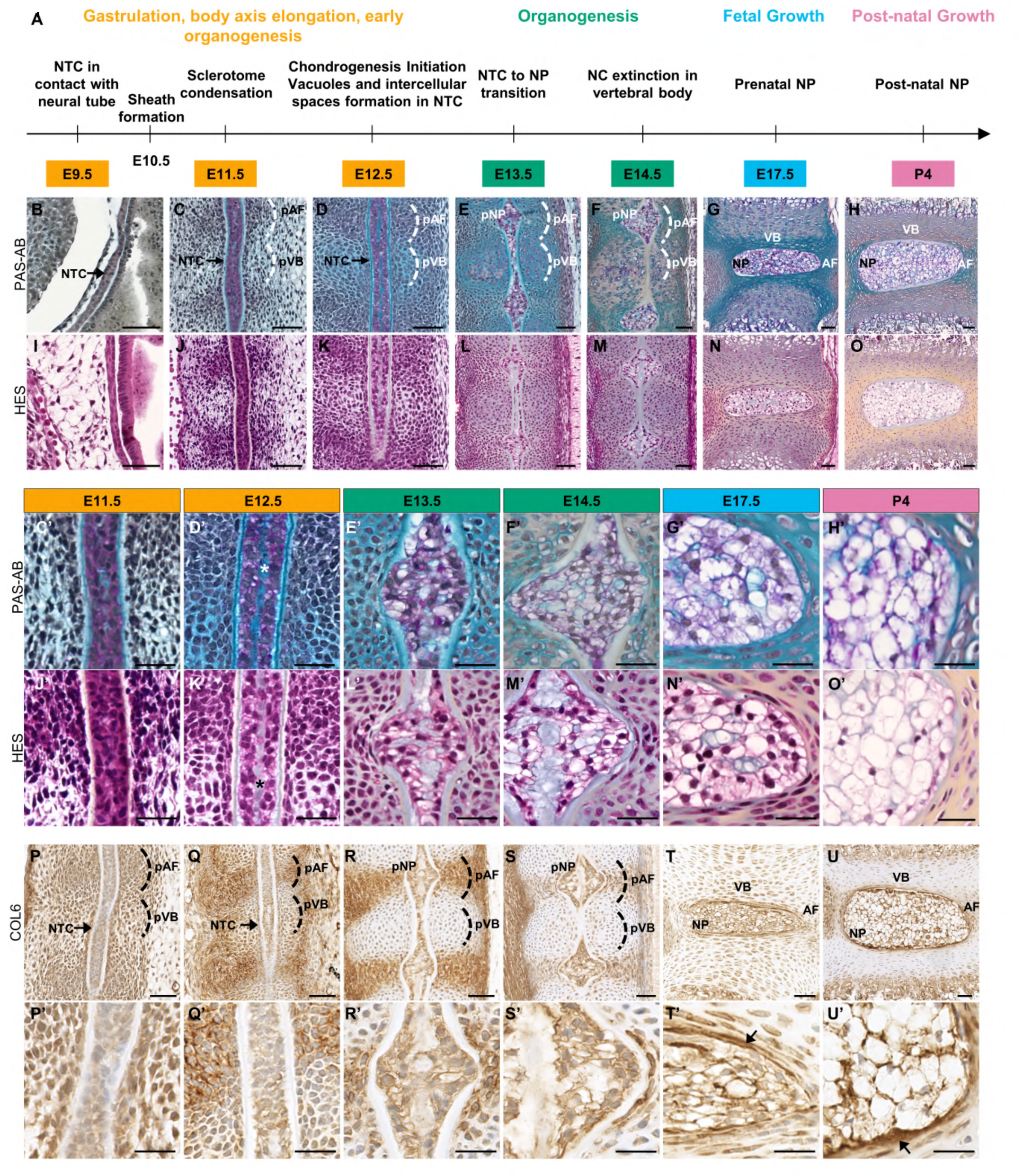
Temporal histological atlas of the notochord and sclerotomes morphological changes during disc development. **(A)**. Schematic summarizing key steps of notochord transformation into the nucleus pulposus during embryonic and postnatal stages. **(B-O)**. Rostro-caudal oriented sections (top-to-bottom) stained with PAS-Alcian Blue (PAS-AB) and Hematoxylin Eosin Saffron (HES) at E9.5 (B, I), E11.5 (C, J), E12.5 (D, K), E13.5 (E, L), E14.5 (F, M), E17.5 (G, N), and P4 (H, O). Note the distinct fusiform NC enlargement at E13.5. Scale bar = 50 µm. (C’–H’; I’–O’) Magnified views of PAS-AB and HES sections. Scale bar= 25 µm. PAS-AB stains nuclei, glycosaminoglycans, and acidophilic structures in blue-purple, blue, and red, respectively. HES stains nuclei, ECM, and cytoplasm in blue-purple, orange, and pink. Stars in D’ and K’ indicate intercellular space at the notochord midline. (P–U) Immunohistochemistry for collagen VI (COL6) at E11.5–P4 shows dynamic matrix deposition. Note the presence of COL6 in the growth plate at P4 (U). (P’, U’) Higher magnifications of COL6 staining. Arrows highlight COL6 deposition at the NP periphery. Scale bar = 25 µm. AF: annulus fibrosus, NP: nucleus pulposus, NTC: notochord, pAF: prospective annulus fibrosus, pNP: prospective nucleus pulposus, pVB: prospective vertebrae, VB: vertebrae.

At E9.5 (20-29 somite pairs) the rod-like notochord structure composed of up to three rows of embryonic notochordal cells (eNCs) in diameter, lies directly beneath the ventral side of the neural tube (Fig. 1B, I; also described in Götz et al., 1995: PMID: 7781042). The formation of the perinotochordal sheath, an acellular laminin- and collagen-rich basement membrane begins around E10.5, concurrently with the migration of the sclerotome mesenchymal cells, which differentiate from the ventral part of the somite, move medially and form the perinotochordal tube around the notochord (data not shown; also described previously^22,23^). At E11.5 (40-45 somite pairs; Fig. 1C, J), eNCs are small, tightly packed within the firm perinotochordal sheath into a rod-like structure approximately 20 µm in diameter. The sclerotomal compartment, initially unsegmented, acquires at this stage a metameric organization with marked regions of different cell density clearly visible (Fig. 1C’, J’). These phenomena of mesenchymal sclerotome condensation or segmentation result in highly condensed areas corresponding to the disc primordium (or anlagen), that will give rise to the inner and outer annulus fibrosus (AF), and less condensed intervening regions of the vertebral primordium, that will undergo cartilaginous differentiation to form the VB^24^. At E12.5 (50 to 55 somite pairs), chondrogenesis initiation can be observed at the level of the presumptive vertebrae, marking the early phase of the formation of cartilage before endochondral ossification, during which loose sclerotome cells differentiate into chondrocyte producing ECM (Fig. 1D). At E12.5, H&E staining reveals the appearance of numerous small vacuoles or vesicles (Fig. 1K, K’) while PAS-AB staining highlights the formation of GAG-rich matrix intercellular spaces (Fig1. D’). Remarkably, a distinct acellular gap filled with ECM is observed in the center of the notochord at this stage (Fig. 1D, D’, K, K’). At E13.5 when somitogenesis ends (up to ∼65 somite pairs), the transition from the notochord to the NP is observed with the accumulation of NCs nuclei at the level of the prospective disc (pAF), while a decrease occurs at the level of prospective VB (pVB; Fig. 1E, L). It has been suggested that the developing VB exerts a mechanical stimulus on the embryonic notochord that induces the relocation of NCs towards the IVD area creating a characteristic pattern of swellings and constrictions resembling a string of beads in shape^22,25,26^. This process is commonly referred as the involution or segmentation of the notochord. At this stage a significant enlargement of the vacuoles is observed in the forming NP which grows to approximately 70 µm in diameter (Fig. 1E’, L’). At E14.5, the NCs have completely disappeared from the area of the forming VB where only basal lamina remained. In contrast, large vacuolated / physaliferous notochordal cells (vNCs) which are distinct from the eNCs, persist and expand at the level of the future disc to shape the newly formed NP (Fig. 1F, M, F’, M’). As the notochord changes to form the NP, PAS-AB staining reveals regions where GAG-rich ECM accumulates, particularly in the future IVD (Fig. 1F, F’). At E17.5 (Fig. 1 G, N, G’, N’) and P4 (Fig. 1H, O, H’, O’), the NP has expanded and adopted a flattened cylindrical shape typical of the adult disc. The outer layers consist of concentric rings of cells and belong to the AF. At these stages, the NP consists primarily of vNCs.

*Col6a1* gene encodes a key ECM molecule whose expression progressively increases in the IVD from development through postnatal growth and into adulthood^27,28^. Immunodetection was conducted to precisely assess COL6 distribution from E11.5 to E17.5 and P4 (Fig. 1P-U). COL6 was detected in the notochord ECM, except in the sheath, from E11.5 onward (Fig. 1P’-S’). At E17.5 and P4, a marked positive deposition of COL6 was observed at the periphery of the formed NP (Fig. 1. T’, U’). At E11.5, diffuse COL6 staining was detected in pericellular regions of the sclerotomes (Fig. 1P, P’). Remarkably, from E12.5 onward, a stronger COL6 signal marked areas of condensed sclerotome and later, the forming AF (Fig. 1 Q’, R-U, R’-U’). This analysis highlighted that this marker delineates the boundaries of pAF throughout maturation, thus changes in COL6 staining can be used to observe the dynamic evolution of the AF structure over time.

Consistently with previous work of Smits and Lefevre, 2003^29^, we observed that cell differentiation and morphogenesis of the notochord proceed in a cranio-caudal sequence. This progression stood out clearly on E14.5 longitudinal section where ongoing morphological changes are apparent (Fig. 2). Indeed, the disc-like shape of the NP at the most advanced thoracic region differs from the morphology of the lumbar region, which is characteristic of newly formed NP, and is even more distinct from the least advanced caudal region showing the typical rod-like morphology.

**Figure 2.**
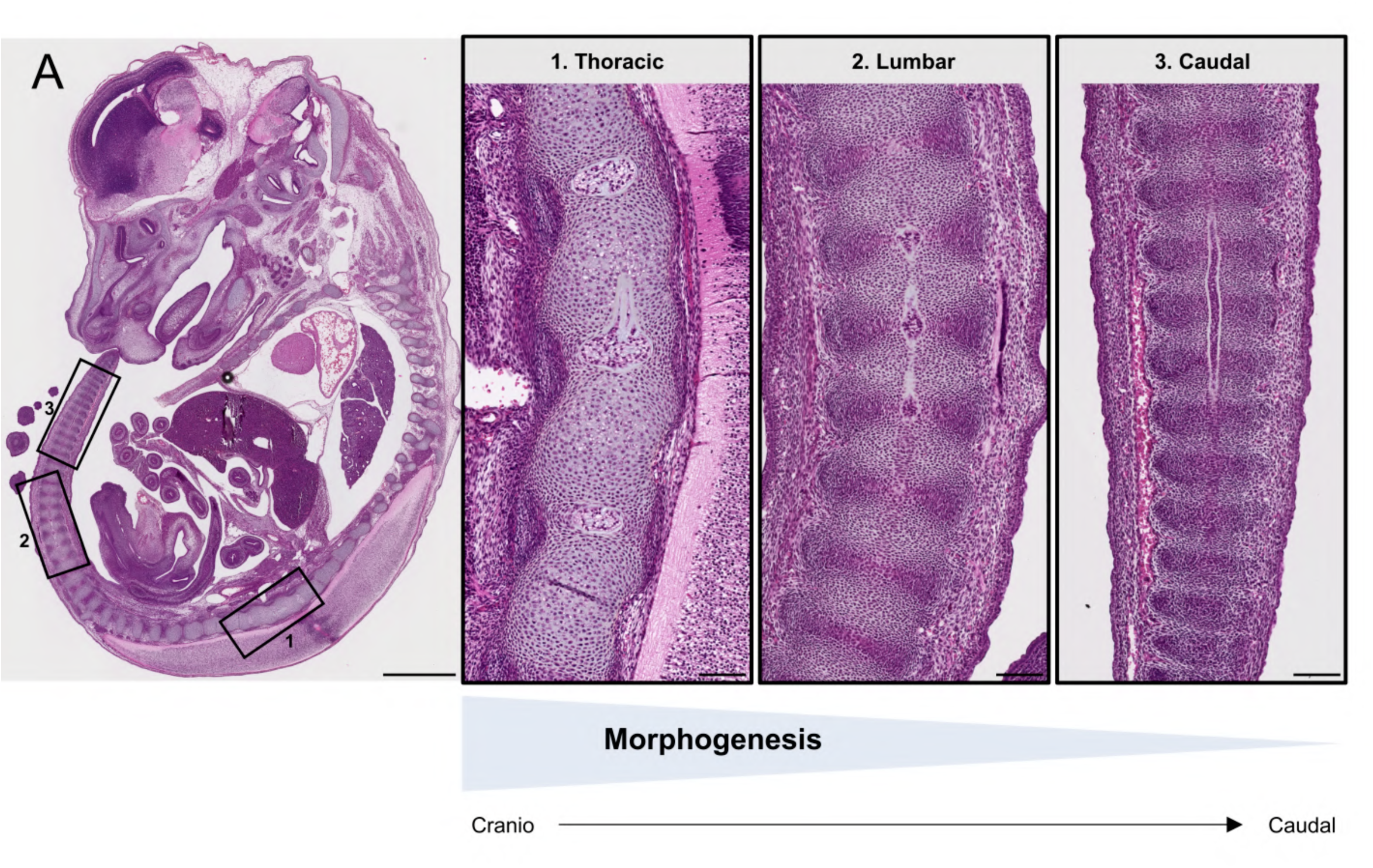
Morphogenesis of the notochord and nucleus pulposus proceed in a cranio-caudal direction. (A). Representative HES staining on mouse histological section at E14.5 and associated schematic representation (B), highlighting distinct morphological features of the notochord on rostro-caudal oriented sections (top-to-bottom). In the thoracic region (box 1) the nucleus pulposus are formed and the notochord is absent in between. In the lumbar region (box 2) the notochord to nucleus pulposus transition is observed with bulging of the notochord where the nucleus pulposus will form and remnant notochord cells where the vertebrae are forming. In the caudal region (box 3), the notochord has still a rod-like shape as classically seen in E12.5 embryo. Scale bar on the whole embryo section = 1mm. Scale bars on the magnified views = 100µm.

### Notochord maturation during nucleus pulposus development is characterized by increased vacuolation and a reduction in cellular density

We observed that the process of NP morphogenesis from the notochordal rod was marked by a striking increase in the volume occupied, and a simultaneous decrease in cellular density. This expansion was also accompanied by ECM accumulation and the appearance of vacuoles at E11.5 (Fig. 1C,C’ and J,J’) that progressively increase in number and size, eventually occupying most of the NP space from E17.5 onward (Fig. 1G,G’). These vacuolar structures, characteristic of notochord-derived cells in the NP, are considered to be related to lysosome-associated organelles^30–35^. As such, LAMP-1 (Lysosomal associated membrane protein 1) was used to monitor vacuole formation and expansion along developmental time.

Confirming histological analyses, LAMP-1 immunohistochemistry identifies the onset of notochordal vacuolation, characterized by the appearance of numerous small intracytoplasmic vacuoles, between E11.5 and E12.5 (Figs. 1J’, K’ and 3A, B, A’, B’). Beginning at E13.5, hypertrophic chondrocytes exhibited vacuole-like structures, that were not further analyzed in this study. Progressive increase in number and enlargement of notochordal intracytoplasmic vacuoles can be observed as eNCs mature into vNCs throughout IVD development (Figs. 1J’-O’ and 3A-F, A’-F’). Interestingly, heterogenous LAMP-1 staining is observed at all stages, with some vacuoles displaying varying staining intensities, suggesting structural differences. To further explore vacuole composition and tissue specificity, co-immunofluorescence of LAMP-1 with another vacuole marker, Caveolin1 (CAV1)^36,37^ and plasma membrane staining were performed at P4 (Fig. 3G, H). LAMP1 and CAV1 were both detected exclusively in NP cells, partially colocalizing. All vacuoles displayed both markers but frequently vacuoles exhibited a predominant staining for either LAMP1 or CAV1. This result confirmed a certain heterogeneity in vacuole composition in the NP as previously reported^38^. To analyse the progression of vacuolation, the LAMP1 positive area, normalised by the number of nuclei, was quantified, based on IHC staining. A consistent increase was demonstrated over time (Table 1; Fig. 3I). While there is only a slight increase between E12.5 and E13.5 (16.03 ± 1.18 to 24.09 ± 1.50, p-value 4.2 x10^-2^), a roughly two-fold increase is observed between E13.5 and E14.5 (up to 50.86 ±2.72, p-value 8.43 x10^-9^), and again between E14.5 and E17.5 (up to 96.1 ± 3.14, p-value 4.55 x10^-15^). In contrast, little difference is seen between E17.5 and P4 (up to 103.11 ± 11.06, p-value 7.66 x10^-1^). In parallel, quantitative analysis of NCs’ nuclei confirmed that cellular density decreased steadily over time from E12.5 to P4 (Table 1; Fig. 3J). Between E12.5 to E14.5, the mean nuclear density decreased from 1.60 ± 0.11 to 0.90 ± 0.04 nuclei/100 μm². The density further reduced by half between E14.5 and E17.5 and between E17.5 and P4, reaching 0.45 ± 0.02 nuclei/100 μm² and of 0.23 ± 0.01 nuclei/100 μm² respectively. The calculated coefficient indicated a strong, negative association between developmental stage and nuclei density.

**Figure 3.**
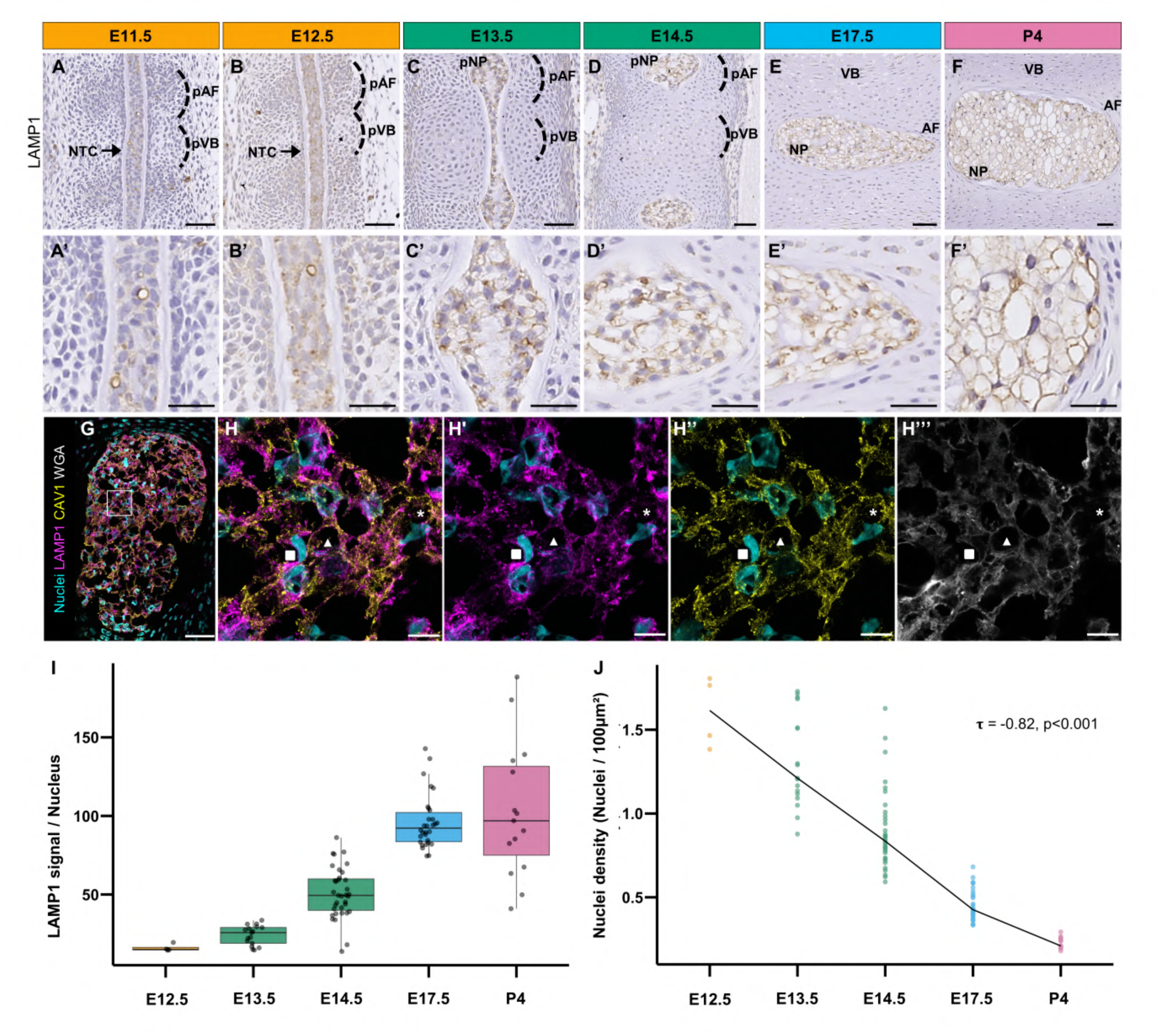
Immunohistological characterization of the intracytoplasmic vacuoles during disc development. **(A-F)** Representative immunohistochemistry of lysosomal-associated membrane protein 1 (LAMP1) on mouse histological sections at E11.5 (A), E12.5 (B), E13.5 (C), E14.5 (D), E17.5 (E), and at P4 (F). Rostro-caudal oriented embryo sections (top-to-bottom). Scale bar = 50µm. **(A’-F’)** Higher magnifications. Scale bar = 25 µm. Nuclei are counterstained with Mayer’s haematoxylin. **(G)** Representative maximum intensity projection images of co-immunofluorescence for LAMP1 and Caveolin-1 on mouse histological sections at P4 with the NP oriented from anterior (left) to posterior (right). Nuclei and membranes were labelled using Hoechst and Wheat Germ Aglutininin (WGA) respectively. Overlays of the dashed box in (G), showing vacuole staining types: ▪ LAMP1-predominant, ▴ dual LAMP1/CAV1, and * CAV1-predominant. Scale = 50 µm (G), 10 µm (H–H’’’). (**I**) Quantification of LAMP1 positive area from E12.5 to P4, normalized by nucleus number per region (NTC or NP). Each dot represents the normalized value of one ROI (NTC or ≥1NP) and middle lines represent median values. **(J)** Quantification of cellular density from E12.5 to P4, measured as the number of nuclei per 100μm² of analyzed area (NTC or NP). Each dot represents the value of one ROI (NTC or ≥1NP). Black line indicates median per stage. Kendall’s test was calculated accross ROI. AF: annulus fibrosus, NP: nucleus pulposus, NTC: notochord, pAF: prospective annulus fibrosus, pVB: prospective vertebrae, VB: vertebrae.

**Table 1.** Quantification of LAMP1-positive area and cellular density across developmental stages from E12.5 to postnatal P4.

| Normalized LAMP1-positive area and cell density |  |  |  |  |  |  |  |  |  |  |
| --- | --- | --- | --- | --- | --- | --- | --- | --- | --- | --- |
| Embryonic stage<br>(Nb of embryos; Nb of sections) | E12.5<br>(1;1) |  | E13.5<br>(2;2) |  | E14.5<br>(4;4) |  | E17.5<br>(3;3) |  | P4<br>(1;2) |  |
| Nb of ROI (NTC or NP ≥1) analyzed | 4 |  | 17 |  | 35 |  | 30 |  | 15 |  |
|  | LAMP1+ | Cellular Density | LAMP1+ | Cellular Density | LAMP1+ | Cellular Density | LAMP1+ | Cellular Density | LAMP1+ | Cellular Density |
| Mean ± s.e.m | 16.03 ± 1.18 | 1.60 ± 0.11 | 24.09 ± 1.50 | 1.31 ± 0.07 | 50.86 ± 2.72 | 0.90 ± 0.04 | 96.13 ± 3.14 | 0.45 ± 0.02 | 103.11 ± 11.06 | 0.23 ± 0.01 |
| Range (min-max) | 14.56–19.53 | 1.38–1.81 | 14.67–33.61 | 0.88–1.73 | 13.75–86.21 | 0.59–1.63 | 74.56–142.79 | 0.33–0.68 | 40.97–188.44 | 0.18–0.29 |

Overall, this analysis suggests that notochord expansion and its transformation into the NP are driven by the phenotypic maturation of eNCs into vNCs. The formation of cytoplasmic vacuoles of increasing volume, along with the deposition of GAG-rich ECM in intercellular space, likely contributes to the observed expansion in tissue volume despite the significant reduction in cellular density as development progresses.

### IVD morphogenesis involves apoptotic events following a cranio-caudal pattern

Changes in cell density and viability have been reported for NC following post-natal growth and during adulthood when they gradually disappear from the NP^39^. Apoptotic cell death has been observed during abnormal disc morphogenesis in mouse genetic models, is prevalent in models of disc degeneration, and is also occurring in both aging and degenerated human discs^11,29,31,32,40–47^. Yet, the description of apoptosis, and its significance in disc morphogenesis during development remain unclear. To bridge this gap of knowledge, we assessed the distribution of programmed cell death along developmental time from E11.5 to E17.5.

First, we performed immunohistochemistry to detect executioner Caspase-3. Positive cells were primarily observed at E11.5 and persisted at E12.5, but were very low or absent at E13.5 and at older E14.5 and E17.5 stages (Fig. 4 A-C and data not shown). Remarkably, Caspase-3 staining was not found in a localized manner in remnant NC at the level of the pVB at E13.5 or E14.5 (Fig. 4C and data not shown). The absence of cell death at these stages indicates that apoptosis does not play a significant role in the elimination of NC at the level of the vertebral disc anlagen during NP formation. In contrast, at earlier stages, apoptotic cells were predominantly observed in the perinotochordal sclerotomes, exhibiting a distinct spatial distribution (Fig. 4A, B). We made the hypothesis that these distinct patterns of cell death, observed at E11.5 and E12.5, could mark the initiation of NP morphogenesis. To test this hypothesis, the number and distribution of Caspase-3+ cells were quantified using a defined analysis scheme (See Experimental Procedures and Supp. Fig. S1A, B). The strategy allowed the measurement of the number of apoptotic cells among notochord or sclerotome across various areas of interest, in particular, distinguishing condensed and non-condensed sclerotomes (alternating formation of pVB and pAF). The quantification also allowed to compare the distribution in the different rostro-caudal regions, i.e., cervical, thoracic, lumbar and/or caudal which could reflect the progression of axial skeleton morphogenesis. Results of the mean and the range of apoptotic cells per categorization are shown in Table 2 and summarized in Fig. 4D at E11.5 and E12.5. No significant difference was found in the mean value of Caspase-3^+^ cells when comparing their distribution in the notochord at pVB and pAF levels. In contrast, Caspase-3^+^ cells were detected in the perinotochordal area with greater number in sclerotome of the pAF area compared to pVB (Fig. 4D). Moreover, apoptotic cells were significantly higher in the sclerotome of pAF area “bordering” the notochord segment versus “anterior” and “posterior” categories. Notably, no apoptotic cell was found in sclerotomes cells in parasagittal sections further away from notochord segment. In a second phase, we examined in more detail the distribution of Caspase-3+ cells in sclerotomes area “bordering” the notochord as well as in notochord itself along the cervical, thoracic, lumbar and caudal regions (Table 3). The results showed that although not significant (Dunn’s test), the number of apoptotic cells per surface area at stage E11.5 was the highest at the caudal region, which at this stage corresponds to the presumptive lumbar and caudal parts together (Table 3). At stage E12.5, no more apoptotic cells were detected in sclerotomes of the cervical region, while only a few were observed in the thoracic region. However, the comparison of the number of apoptotic cells per μm² using Dunn’s test showed that apoptosis was statistically higher in the lumbar and caudal regions when comparing the number of apoptotic events across the different rostro-caudal regions. In summary, apoptotic events were observed at both stages, with the overall incidence of apoptosis decreasing across all regions from E11.5 to E12.5 and with distinct variation in apoptosis levels along the rostro-caudal axis. Thus, the quantification revealed that the degree of apoptosis in sclerotome cells near the notochord fragment increased progressively from head to tail as shown in Fig. 4E. This progression of cell death in perinotochordal cells was even more marked at E12.5 with statistical differences between regions. This particular distribution, and the variation between stages E11.5 and E12.5, support the hypothesis that a cranio-caudal wave of apoptotic events in perinotochordal sclerotomes may contribute to the morphogenetic process of NP formation from the notochord, which proceeds in a similar head-to-tail sequence (Fig. 2).

**Figure 4.**
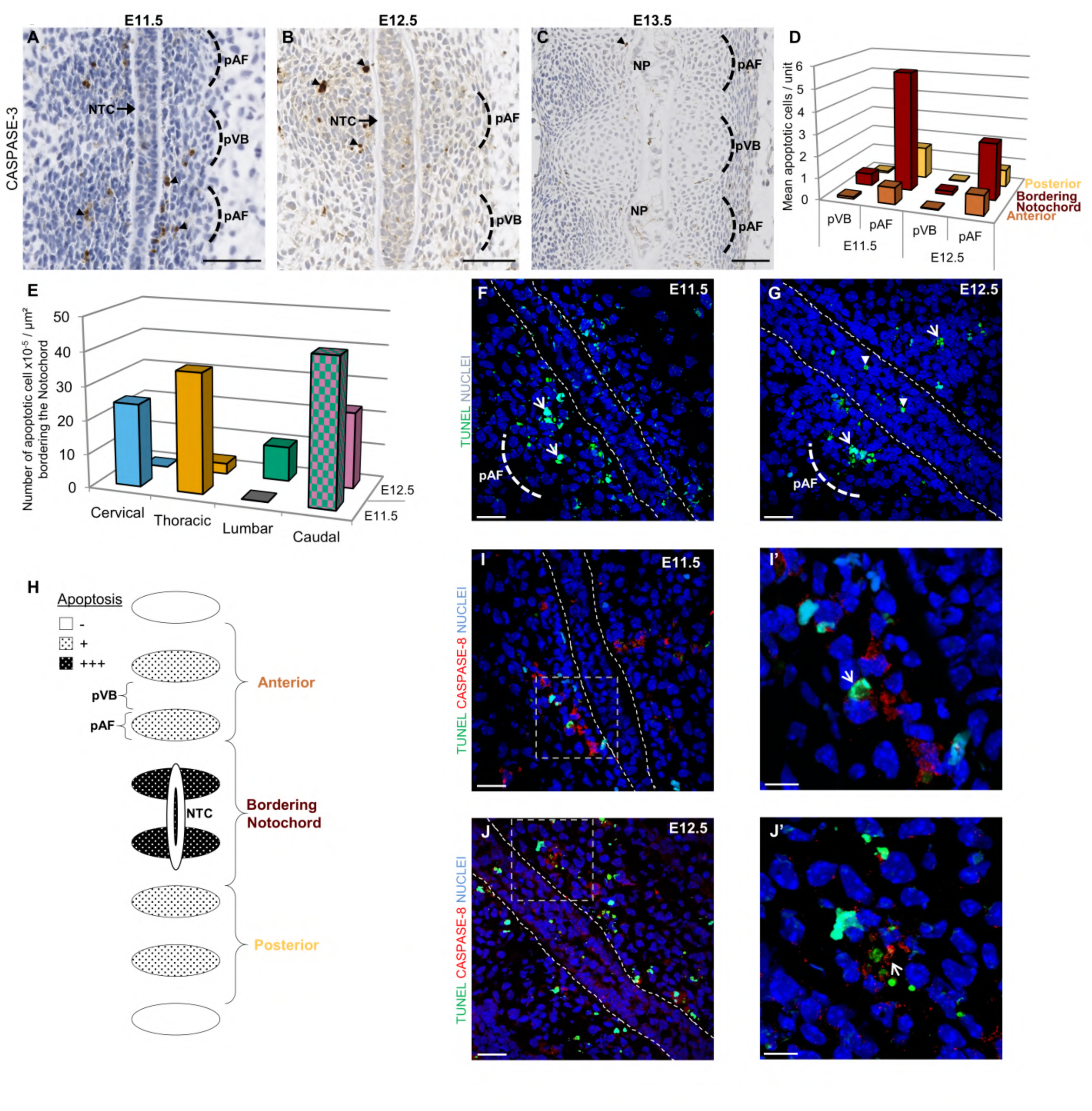
Apoptosis contributes to IVD morphogenetic changes following a cranio-caudal sequence. **(A-C)** Caspase-3 immunohistochemistry at E11.5, E12.5, and E13.5 on rostro-caudal oriented mouse embryo sections (top-to-bottom). Arrowheads mark Caspase-3⁺ cells. Scale bars = 50 µm. **(D)** Quantification of mean apoptotic cells per unit at pVB and pAF levels for E11.5 and E12.5. **(E)** Quantification of number of apoptotic cells per µm² in perinotochordal regions across cervical, thoracic, lumbar, and caudal levels. At E11.5, lumbar and caudal sclerotomes are not yet condensed, thus values reflect total caudal sclerotome area. At E12.5, sclerotome cells of the pAF are condensed at all axial levels. **(F-G)** Representative TUNEL staining at E11.5 (F), and E12.5 (G). White arrow heads indicate TUNEL^+^ cells. Embryo sections are oriented from anterior (left) to posterior (right). Dotted line highlights the notochord. Scale bar = 25µm. **(H-I)** Representative immunofluorescence staining of TUNEL and Caspase-8 at E11.5 and E12.5. White arrow heads indicate TUNEL^+^/Caspase-8^+^ cells. Scale bar = 25µm. **(H’-I’)** Higher magnification of dashed boxes. Scale bar = 10 µm. **(J)** Schematic illustrating the spatial distribution of apoptotic cells as determined by Caspase-3 quantification. NP: nucleus pulposus, NTC: notochord, pAF: prospective annulus fibrosus, pVB: prospective vertebrae.

**Table 2.** Distribution of Caspase-3-positive cells in pVB, pAF and notochord along the whole axis. Dunn’s (1964) test of multiple comparisons following a significant Kruskal-Wallis test with a correction to control the experiment-wise error rate were applied to the quantification data to compare pAF with pVB and NTC and Border. with Ant. and Post. categories. P-values adjusted with the Holm method were ***p< 0.001. # indicate the total mean value for NTC as no significant difference is observed when comparing Caspase-3^+^ cells distribution in the notochord between pVB and pAF levels. Ant. = pVB and pAF positioned “anterior” to the notochord segment; Border. = pVB and pAF bordering the notochord; Post. = pVB and pAF positioned “posterior” to the notochord segment.

| Caspase 3+ cells in pVB, pAF and notochord along the whole axis |  |  |  |  |  |  |  |  |  |  |  |  |  |  |
| --- | --- | --- | --- | --- | --- | --- | --- | --- | --- | --- | --- | --- | --- | --- |
| Embryonic stage<br>(Nb of embryos;<br>Nb of sections) | E11.5<br>(4;47) |  |  |  |  |  |  | E12.5<br>(4;55) |  |  |  |  |  |  |
|  | Sclerotome cells |  |  |  |  |  | Notochord<br>cells | Sclerotome cells |  |  |  |  |  | Notochord<br>cells |
|  | pVB<br>(n=329) |  |  | ***pAF<br>(n=276) |  |  | NC | pVB<br>(n=619) |  |  | ***pAF<br>(n=542) |  |  | NC |
|  | Ant.<br>(n=96) | ***Border<br>(n=137) | Post.<br>(n=96) | Ant.<br>(n=77) | ***Border<br>(n=129) | Post.<br>(n=70) | Total #<br>(n=142) | Ant.<br>(n=153) | ***Border<br>(n=305) | Post.<br>(n=161) | Ant.<br>(n=127) | ***Border<br>(n=282) | Post.<br>(n=133) | Total #<br>(n=382) |
| Mean ± s.e.m | 0.09± 0.09 | 0.55± 0.19 | 0.08± 0.07 | 0.77± 0.37 | 5.49± 0.92 | 1.47± 0.45 | 0.08 ± 0.04 | 0.03± 0.01 | 0.16± 0.06 | 0.03± 0.01 | 0.90± 0.25 | 2.61± 0.64 | 0.78± 0.21 | 0.09 ± 0.04 |
| Range<br>(Min-Max) | 0.00-0.50 | 0.00-1.27 | 0.00-0.50 | 0.21-2.67 | 1.33-8.20 | 1.11-4.0 | 0.00-0.26 | 0.00-0.11 | 0.00-0.79 | 0.00-0.17 | 0.00-3.25 | 0.00-7.90 | 0.00-2.33 | 0.00-0.52 |

**Table 3.**
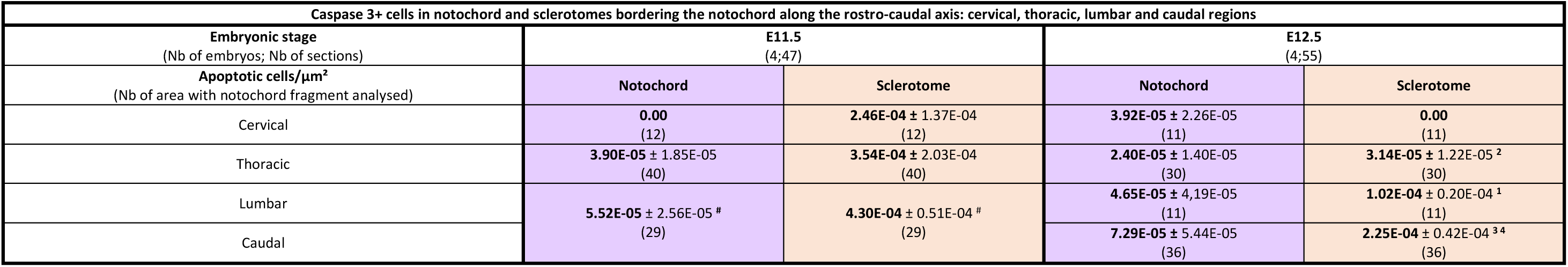
Distribution of Caspase-3-positive cells in sclerotomes bordering the notochord along the rostro-caudal axis: cervical, thoracic, lumbar and caudal regions. Dunn’s test (1964) for multiple comparisons was applied to the quantification data following a significant Kruskal-Wallis test with a correction to control the experiment-wise error rate. No statistic difference were found between regions except at E12.5 in sclerotomes cells between cervical and lumbar ^1^ (p< 0.01), thoracic and lumbar ^2^ (p< 0.05), thoracic and caudal ^3^ (p< 0.001) and cervical and caudal ^4^ (p< 0.001). P-values adjusted with the Holm method. # indicate the number of apoptotic cells relative to the total area of sclerotomes analysed around the notochord segment in presumptive lumbar and caudal parts of E11.5 embryos as at this stage the sclerotomes of the pAF are not condensed yet.

Apoptotic cell death was also detected by TUNEL staining and was consistent with results described for Caspase-3 immunostaining (Fig. 4F, G, for E11.5 and E12.5, and data not shown for E13.5). Remarkably, confocal microscopy with optical sectioning revealed a high level of cell death at E12.5 localized at the midline of the notochord along the entire rostro-caudal axis, creating a continuous median intercellular gap in its rod-like structure (Fig. 4G and Supp. Fig. S1C for E12.5 TUNEL assay). Note that, contrasting with the very low cell death detected in the forming axial skeleton from E13.5, extensive cell death remains visible in the most caudal part of the notochord in E13.5 embryos, where its morphology still retains the rod-like aspect characteristic of the E12.5 stage (Supp. Fig. S1D for caudal part of E13.5 TUNEL assay). Cell death was also clearly observed on embryonic sections stained with both PAS-AB and H&E where notochord exhibits numerous pyknotic nucleus and cytoplasmic eosinophilia suggestive of apoptosis (Supp. Fig. S1E, F). Altogether, this analysis reveals specific patterns and a temporal shift of cell death within a short window from E11.5 to E12.5. This occurs in the sclerotome localized in pAF areas adjacent to the notochord. During the same periods, the notochord itself exhibits a pronounced level of cell death along the midline (summarized in Fig. 4H).

Lastly, cell death was assessed by TUNEL in combination with immunofluorescence for the apical initiator Caspase-8, a key mediator of the extrinsic pathway, to determine whether apoptosis in this developmental context is triggered through autocrine or paracrine interactions. The extrinsic pathway that initiates apoptosis is induced by death ligand binding to a death receptor at the cell surface. Of particular interest, Caspase-8 signal was detected exclusively in sclerotome cells of the pAF areas adjacent to notochord segment and was found localized in area of TUNEL positive cells along the rostro-caudal axis of E11.5 and E12.5 embryos, indicating transient activation of apoptosis through the death receptor pathway (Fig. 4I, I’, J, J’).

### NC proliferation dynamics follows a cranio-caudal pattern during IVD morphogenesis

Although our results shows that cell death likely plays a role in IVD morphogenesis, it alone cannot account for segmental expansion and constriction of the NCs. Therefore, we investigated whether differential cell proliferation also contributes to this morphogenetic process. First, we assessed the distribution of mitotic cells by performing anti-phospho-histone H3 antibody (pHH3) immunohistochemistry along developmental time^48^. Remarkably, intensely stained replicating cells appeared similarly distributed in notochordal segments at pAF and pVB levels from E11.5 to E14.5 (Fig. 5A-D). Indeed, qualitative examination of the sections revealed no distinct pattern of actively dividing cells across the embryonic stages analysed. These results indicate that pHH3^+^ nuclei do not exhibit clustering beyond what would be expected from a random distribution. In particular, even though at E13.5, the reduced cellularity is occurring in the constricted areas of the notochord, replicating pHH3^+^ NCs were still detectable at pVB level (Fig. 5C). Based on the overall distribution of the mitotic cell marker in notochord segments, we concluded that differential cell proliferation could not account for the segmental expansion of the NCs at the pAF levels, which begins to emerge at E13.5.

**Figure 5.**
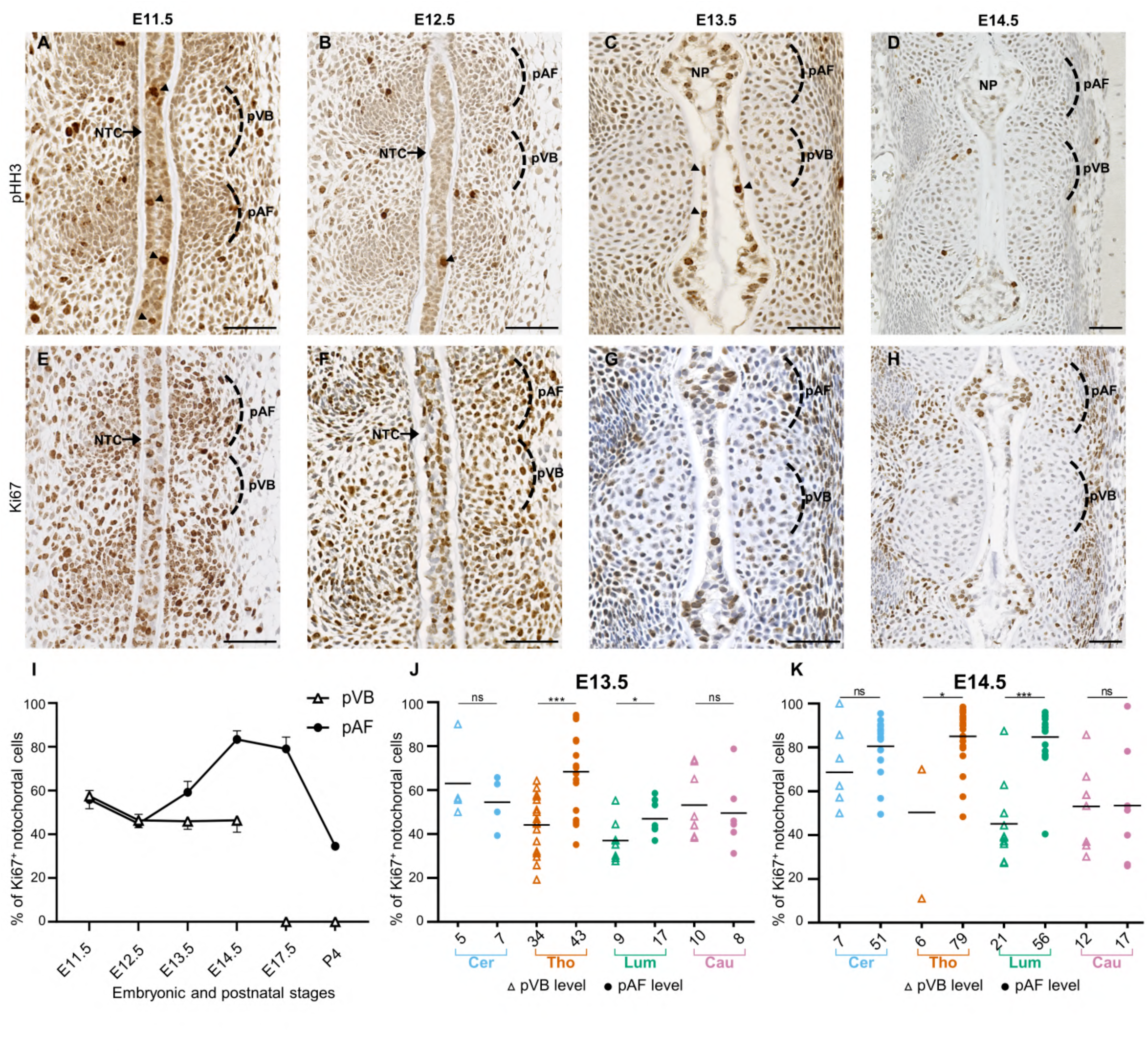
Proliferative capacity of notochordal cells contributes to IVD morphogenetic changes following a cranio-caudal sequence. **(A-D)** Representative immunohistochemistry of phosphohistone 3 (pHH3) on mouse histological sections at E11.5 (A), (N= 3 embryos; n= 6 sections), E12.5 (B), (N= 2 embryos; n= 4 sections), E13.5 (C), (N= 7 embryos; n= 14 sections), and E14.5 (D), (N= 3 embryos; n= 9 sections). **(E-H)** Representative immunohistochemistry of Ki-67 on mouse histological sections at E11.5 (E), E12.5 (F), E13.5 (G) and E14.5 (H). **(I)** Quantification of Ki-67^+^ notochordal cells according their proximity with the pAF or PVB along developmental time. Note: after E14.5 the intervertebral disc is formed and the notochord is absent from the area where the vertebrae is formed. **(J-K)** Quantification of Ki-67⁺ notochordal cells at E13.5 (J) and E14.5 (K) across rostro-caudal levels (cervical, thoracic, lumbar, caudal), and at the proximity to pAF or pVB. The x-axis indicates number of sections analysed. Statistical analysis: Kruskal-Wallis test followed by Dunn’s post hoc test (1964) with Holm correction for multiple comparisons. Asterisks indicate significance (*p < 0.05; ***p < 0.001). Embryo sections are displayed in a rostro-caudal orientation (top-to-bottom). Scale bar = 50µm. NP: nucleus pulposus, ns: non-significant, NTC: notochord, pAF: prospective annulus fibrosus, pVB: prospective vertebrae.

To further investigate the overall proliferative potential of the NCs during IVD morphogenesis, we performed immunohistochemistry for Ki-67 to label proliferative cells (Fig. 5E-H). Interestingly, we observed that as early as E11.5, only a subset of the notochord population was stained, indicating the remaining cells had entered a quiescent (G0) state. Moreover, from E13.5 stage, variations in the distribution of the Ki-67^+^ proliferative cell subpopulation became evident. We hypothesized that these distinct patterns of proliferation activity may contribute to NP morphogenesis. To test this hypothesis, the number and distribution of Ki-67^+^ cells were quantified using a defined analysis scheme. The strategy allowed us to count proliferative NCs while distinguishing them into two categories-“at the level of pVB” or “at the level of pAF”- and to compare their distribution across the four different rostro-caudal regions (Supp. Fig. S1G). In the early stages, the distribution of Ki-67^+^ cell in the notochord was homogeneous between pVB and pAF categories, with above 50% and 40% of Ki-67^+^ NCs detected at E11.5 and E12.5 respectively, regardless of the rostro-caudal regions (Fig. 5I and Table 4). In contrast, during the transition from notochord to NP at E13.5 and E14.5 stages, the percentage of Ki-67^+^ cells progressively increased in notochord segments at the level of pAF compared to the pVB. A difference of 13% and 37% of proliferative cells was observed in pAF areas at E13.5 and E14.5, respectively (Fig. 5I and Table 4). At E14.5 stage, the Ki-67^+^ count in pAF was 1.8 times higher than in pVB across the entire rostro-caudal axis, suggesting that NCs in the forming disc preferentially retain their proliferative capacity. From E14.5 onward, no remnant NC in the developing VB is observed as the NP is formed. Within the NP structure, the overall proportion of proliferating NCs progressively decreased across the E14.5, E17.5, and P4 stages, with a sharp decline from approximatively 80% to 35% after birth (Fig. 5I; Table 4). We then examined in greater detail the proliferative capacity of NCs across distinct axial regions, with a specific focus on the transition from notochord to NP at E13.5 and E14.5 (Fig. 5G, H ; Table 5). This analysis revealed that the proliferative capacity of NCs at the pAF level was higher than at the pVB level, particularly at the thoracic and lumbar regions. At E13.5, proliferation was statistically more pronounced in the thoracic region compared to lumbar whereas at E14.5, it became statistically more prominent in the lumbar region. No statistical difference was observed in the caudal regions, where the NP has not yet formed at these stages, and the notochord retains a comparable morphology to E12.5 (Fig. 2).

**Table 4.** Quantification of Ki-67-positive notochordal cells according to the proximity with the prospective annulus fibrosus or prospective vertebrae.

| Percentage of Ki67+ cells in notochord and nucleus pulposus localised at the pVB and pAF levels from E11.5 to P4 |  |  |  |  |  |  |  |  |  |  |  |  |
| --- | --- | --- | --- | --- | --- | --- | --- | --- | --- | --- | --- | --- |
| Embryonic Stage<br>(Nb of embryos;<br>Nb of sections) | E11.5<br>(3;20) |  | E12.5<br>(3;7) |  | E13.5<br>(3;11) |  | E14.5<br>(6;21) |  | E17.5<br>(3;13) |  | P4<br>(3;6) |  |
|  | Notochord |  | Notochord |  | Notochord |  | Notochord |  | Nucleus pulposus |  | Nucleus pulposus |  |
|  | pVB | pAF | pVB | pAF | pVB | pAF | pVB | pAF | pVB | pAF | pVB | pAF |
| Nb of unit with notochord<br>fragment analyzed | 7 | 8 | 9 | 10 | 12 | 12 | 14 | 17 | NA | 3 | NA | 49 |
| % Ki67+ cells ± s.e.m | 57.43 ± 5.69 | 55.90 ± 4.14 | 46.35 ± 3.32 | 44.91 ± 4.25 | 45.95 ± 3.68 | 59.25 ± 4.99 | 46.39 ± 5.39 | 83.35 ± 3.91 | NA | 79.04 ± 5.44 | NA | 34.54 ± 1.72 |

**Table 5.** Quantification of Ki-67-positive notochordal cells localised at the pVB or pAF level along the cervical, thoracic, lumbar and caudal regions. Only E13.5 and E14.5 embryonic stages show statistically significant differences. Dunn’s test (1964) for multiple comparisons was applied to the quantification data following a significant Kruskal-Wallis test with a correction to control the experiment-wise error rate. No statistic difference was found between pAF and pVB at E11.5 and E12.5. At E13.5, a higher number of Ki-67^+^ cells is found at the pAF level in the thoracic (***p< 0.001) and lumbar (*p< 0.05) regions. At E14.5, a higher number of Ki-67^+^ cells is found at the pAF level in the thoracic (*p< 0.05) and lumbar (***p< 0.001) regions. P-values adjusted with the Holm method.

| Number of Ki67 + cells in notochord localised at the pVB or pAF level along the cervical, thoracic, lumbar and caudal regions |  |  |  |  |  |  |  |  |
| --- | --- | --- | --- | --- | --- | --- | --- | --- |
| Embryonic Stage<br>(Nb of embryos; Nb of sections) | E11.5<br>(3;35) |  | E12.5<br>(3;14) |  | E13.5<br>(3;38) |  | E14.5<br>(6;31) |  |
|  | Notochord |  | Notochord |  | Notochord |  | Notochord |  |
|  | pVB | pAF | pVB | pAF | pVB | pAF | pVB | pAF |
| **Cervical** |  |  |  |  |  |  |  |  |
| (Nb of unit with NTC fragment analyzed) | (20) | (21) | (11) | (12) | (5) | (7) | (7) | (51) |
| % Ki67+ cells ± s.e.m | 55.53 ± 9.67 | 54.15 ± 7.82 | 42.36 ± 7.97 | 41.76 ± 11.26 | 61.86 ± 7.58 | 59.03 ± 9.54 | 62.79 ± 18.20 | 77.67 ± 6.44 |
| Total nuclei per unit<br>Mean ± s.e.m | 25.21 ± 1.79 | 31.00 ± 1.67 | 27.64 ± 6.56 | 35.42 ± 3.83 | 21.08 ± 8.42 | 31.70 ± 1.70 | 6.87 ± 0.70 | 30.95 ± 2.55 |
| **Thoracic** |  |  |  |  |  |  |  |  |
| (Nb of unit with NTC fragment analyzed) | (33) | (36) | (18) | (20) | (34) | (43) | (6) | (79) |
| % Ki67+ cells ± s.e.m | 52.40 ± 7.21 | 48.24 ± 11.28 | 47.49 ± 6.02 | 51.91 ± 5.76 | 43.33 ± 3.63 | 66.86 ± 9.72 *** | 51.72 ± 19.63 | 87.26 ± 3.38 * |
| Total nuclei per unit<br>Mean ± s.e.m | 16.20 ± 11.70 | 24.03 ± 3.36 | 46.58 ± 10.51 | 39.76 ± 6.54 | 18.39 ± 1.45 | 24.88 ± 0.64 | 5.78 ± 1.68 | 33.87 ± 2.97 |
| **Lumbar** |  |  |  |  |  |  |  |  |
| (Nb of unit with NTC fragment analyzed) | (13) | (14) | (3) | (5) | (9) | (17) | (21) | (56) |
| % Ki67+ cells ± s.e.m | 77.57 ± 11.83 | 72.31 ± 3.43 | 48.67 ± 0.20 | 45.65 ± 4.82 | 35.77 ± 0.12 | 49.80 ± 3.33 * | 42.00 ± 6.66 | 86.12 ± 4.47 *** |
| Total nuclei per unit<br>Mean ± s.e.m | 17.51 ± 5.41 | 30.54 ± 3.82 | 53.50 ± 10.50 | 52.00 ± 16.00 | 45.45 ± 1.95 | 28.86 ± 2.69 | 13.78 ± 2.63 | 44.65 ± 3.81 |
| **Caudal** |  |  |  |  |  |  |  |  |
| (Nb of unit with NTC fragment analyzed) | NA | NA | (2) | (7) | (10) | (8) | (12) | (17) |
| % Ki67+ cells ± s.e.m | NA | NA | 50.00 | 34.66 ± 11.14 | 55.18 ± 8.87 | 49.71 ± 12.51 | 48.57 ± 8.70 | 67.58 ± 8.58 |
| Total nuclei per unit<br>Mean ± s.e.m | | | 24.00 ± 0.00 | 51.04 ± 5.29 | 49.06 ± 2.94 | 42.50 ± 4.25 | 20.81 ± 5.07 | 22.93 ± 4.12 |
| % Ki67+ cells all regions ± s.e.m | 57.43 ± 5.69 | 55.90 ± 4.14 | 46.35 ± 3.32 | 44.91 ± 4.25 | 45.95 ± 3.19 | 59.25 ± 4.32 | 46.39 ± 5.39 | 83.35 ± 3.91 |

Our results provide evidence that Ki-67 expression distinguishes patterns of proliferating NCs that collectively contribute to NP formation. They indicate a dynamic shift in Ki-67^+^ cells distribution, aligning with the cranio-caudal morphogenesis process. They also reveal differences in NCs proliferative capacities between regions of the forming NP and vertebrae. Altogether our findings support the hypothesis that the spatial and temporal patterns of proliferation within the notochord contribute to the morphogenetic process of NP formation.

## Discussion

Morphogenesis involves diverse processes that shape embryonic tissues and structures at the origins of the organs. It can involve cell proliferation, apoptosis, migration, rearrangement, cellular orientation, adhesion, and mechanical forces. Here, we highlight tightly linked events of controlled proliferation and apoptosis coordinated during mouse spine development, that contributes to notochord expansion and NP tissue specialization in the forming IVD. We showed that significant differences in proliferation and apoptosis at specific developmental stages contribute to normal progression of NP morphogenesis by causing irreversible changes in notochord and sclerotome cells arrangements. Additionally, we showed that these processes are accompanied by the phenotypic maturation of eNCs into mature vNC, marked by prominent cytoplasmic vacuoles and ECM deposition, therefore sustaining NP morphogenesis despite progressive decline in cellular density. Our work provides a precise description and quantitative assessments of these specific morphogenetic events, establishing a reference resource for phenotypic analysis in IVD development, in particular in NP morphogenesis, and enabling cross-study comparisons in mice.

During the early developmental phase (E11.5-E12.5; summarized in Fig. 6), our analyses revealed a transient activation of apoptosis via the death receptor pathway in the perinotochordal sclerotome, preferentially localized at the level of the forming disc. Quantitative analysis confirmed that cell death follows a temporal cranio-caudal shift. Apoptosis is a highly regulated form of programmed cell death that classically involves the intrinsic (mitochondrial-dependent) and extrinsic (death receptor or Fas-dependent) pathways. Various works have shown that caspases play key roles in controlling cellular shape et tissue remodelling^49^. The apical Caspase-8 which initiates programmed cell death induced by the extrinsic pathway was detected exclusively, in sclerotome cells of the pAF at the proximity of the notochord, following a cranio-caudal sequence at E11.5 and E12.5. One plausible explanation for the localized occurrence of apoptosis in the condensed perinotochordal sclerotome is that it creates space potentially alleviating physical constraints, thereby facilitating the accumulation of proliferative vacuolated NCs which collectively contribute to the formation of the NP. Apoptosis within the condensed sclerotome may be determined by signaling cues involving SHH, WNT or TGF-β pathways known to be involved in apoptotic process. Although the origin of the apoptotic stimulus, whether from the notochord or the sclerotomes, remains to be determined, the results suggest that specific spatial cues, such as changes in the extracellular environment, play a critical role in determining where and when a cell undergoes apoptosis. Alternatively, apoptosis between E11.5 and E12.5 may be part of the morphogenetic events promoting cartilaginous inner AF development and possibly eliminating cells with an inappropriate VB or fibrous annulus developmental fate. Our study also highlights that extensive cell death also occurs simultaneously at the midline of the notochord along the entire rostro-caudal axis at E12.5. These apoptotic events appear to lead to the formation of an acellular median gap with no apparent epithelial lining. Concomitantly, with the initiation of vacuolation within the notochord starting from E11.5 along the entire axis, the earliest signs of ECM deposition appear within the newly formed intercellular gap (Fig. 6). While apoptosis is likely to play a role in the transformation of the rod-like notochord into a hollowed structure, the precise function of this process in normal morphogenesis remains to be elucidated. One possibility is that the physical stress associated with vacuole formation activates the intrinsic apoptosis pathway, leading to inner gap formation. This might be essential for creating space for ECM deposition and remodelling, thereby facilitating cellular rearrangement and phenotypic changes during the notochord-to-NP transition^50^.

**Figure 6.**
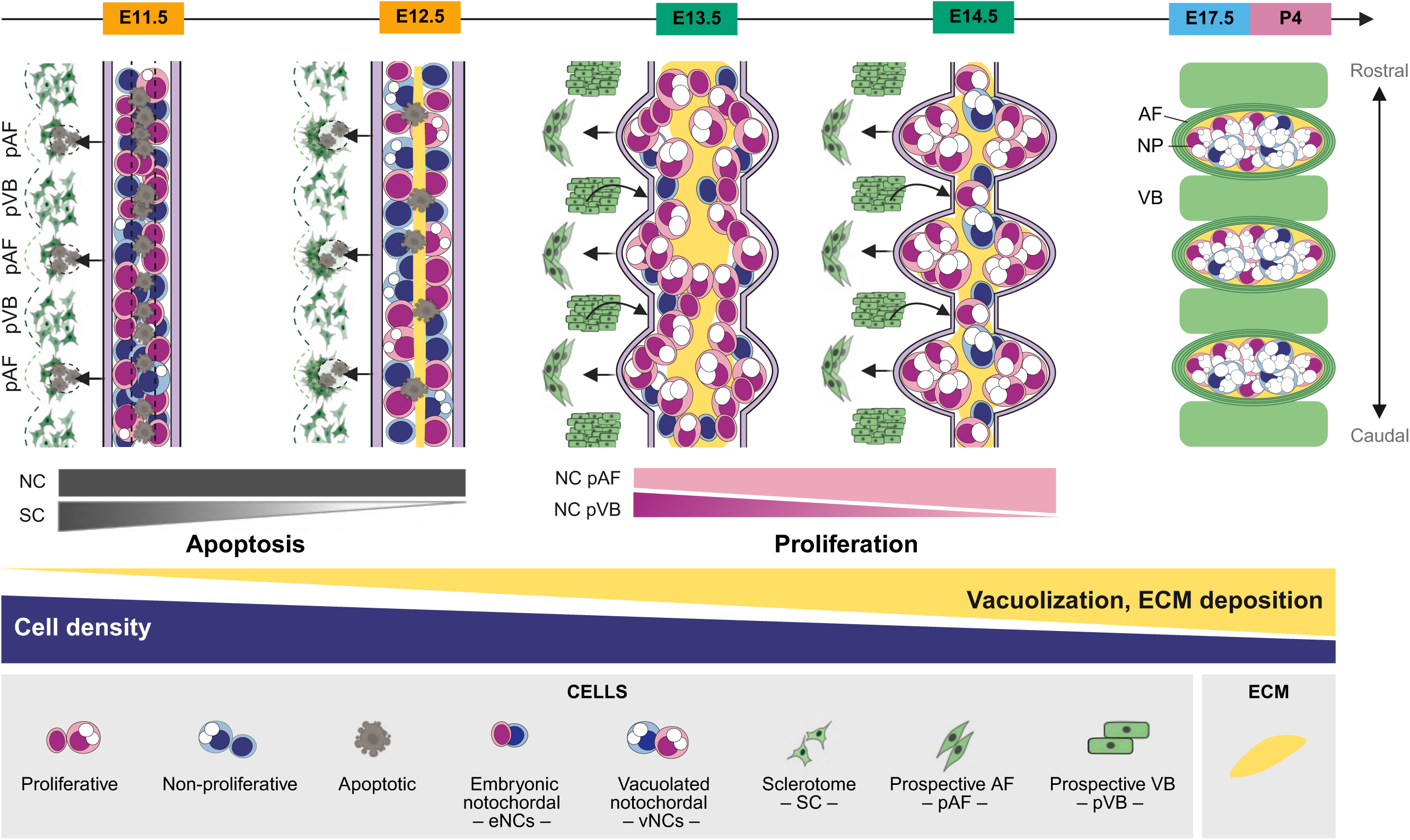
Disc development model. Schematic overview of IVD morphogenesis from E11.5 to E17.5–P4. Major structural changes occur as the NCs accumulate at the future disc sites to form the NP, but are not maintained where the VBs form. Our findings provide evidence for two distinct developmental patterns of morphogenetic events, as illustrated here by thoracic and lumbar levels (see Supp. Fig. S2 for full rostro-caudal axis). The first pattern (E11.5-12.5) involves a synchronous decrease in cell density, an increase in cytoplasmic vacuole number and size, and ECM deposition over time along the axis. Note the early NC apoptosis that uniformly creates a midline gap (white dash) most likely facilitating ECM deposition (yellow). The second pattern (E13.5-14.5) reflects a rostro-caudal progression similar to the one observed in NP morphology with differential apoptosis in sclerotomes (E11.5–12.5) and differential proliferative capacities in NCs located at pAF and pVB levels. Arrows indicate proposed physical cues from pAF apoptosis (straight) and pressure constraint from forming pVB (curved), both guiding NC expansion into pAF, along with the increased ability for NCs to divide in this area. By P4, proliferative NCs remained localized at the NP periphery (Paillat et al., 2023: 10.22203/eCM.v045a06; Zhang et al, 2024: 10.1096/fj.202301217R; Tan et al., 2024: 10.1016/j.celrep.2024.114342). ECM: extracellular matrix; pAF and pVB: prospective annulus fibrosus and vertebrae. Created in https://BioRender.com.

Over the late developmental phase (E13.5-E14.5), results indicate that following apoptosis, distinct spatial and temporal patterns of proliferation contribute to localized tissue expansion and differentiation. Quantitative analyses revealed that proliferative cell distribution within the notochord shifts dynamically, supporting cranio-caudal NP morphogenesis. We hypothesize that earlier cranio-caudal apoptotic wave in sclerotomes could facilitate NP expansion from the notochord which proceed in a similar head-to-tail sequence (Figs. 6 and S2). Our findings suggest that the dynamic interplay between these cellular processes together with progressive increase in ECM deposition and intracytoplasmic vacuoles formation and expansion, as well as cell rearrangement, plays a crucial role in continuously shaping the NP along the rostro-caudal axis of the embryo (Fig. 6). Consistent with previous reports on proliferative characteristics of cells of the ventral node and the derived notochordal plate using BrdU labelling from E7.5 to E14.5^25,51–53^, we showed here that a significant portion of NCs are quiescent from E9.5 onward. A recently published study using EdU injection to map proliferation showed that the enlargement of the NP involves localized changes in notochord proliferation, prior to hypertrophic differentiation of the VB region as assessed by Collagen 10 marker^54^. Supporting this result, our work reveals that NCs proliferative capacity appeared not uniform, and suggests regulatory influences on proliferation at later phases during disc formation (E13.5-E14.5), when the notochord transitions into the NP. In addition, quantitative analyses revealed that the distribution of proliferative NCs shifts dynamically in a cranio-caudal direction, likely contributing to notochord-to-NP morphogenesis through progressive cell expansion and rearrangement. Thus, we hypothesize that distinct proliferative dynamics driven by local signaling occur in specific areas to shape the NP along the rostro-caudal axis (Figs 6 and S2).

Both biochemical and biomechanical signals could be influencing these distinct proliferative behaviors, this remain to be elucidated. Extensive crosstalk between biochemical and mechanical stimuli during tissues morphogenesis and growth is now well established^55,56^. Changes in mechanical force distribution may create differences that influence cell proliferation during subsequent development and morphogenesis of the disc. These could explain that cells located in areas of forming VBs sense forces, in contrast to cell in areas of forming NPs, leading to distinct proliferation behaviors. A specific phenotype described as “rod-like notochord persistence” has been reported by several studies, supporting the idea that physical mechanism may be responsible for the notochord to NP transition, forming the basis of the “pressure” model^9,21,25,57,58^. Mutations affecting *Pax-1* gene expression in the sclerotome results in a higher proliferation rate of NCs at E12.5 and “notochord persistence” at the level of the vertebrae^5,59^. Although the mechanism has not been elucidated, this phenotype suggests that sclerotomes signal back to the notochord to control its expansion strictly at the level of the forming disc. Alternatively, the Repulsion/Attraction model has been suggested, where locally expressed attractant/repulsive signals yet to be determined, could explain notochord cell movement toward the forming NP and/or away of the forming VB^21^. Evidence that NCs migrate or are rearranged under any type of pressure or discrete biochemical signal remains to be determined during disc formation. Further investigations are needed to demonstrate that part of these signals or mechanisms exert influence on the proliferative behavior on the notochord cell as well as on apoptosis patterning within the perinotochordal sclerotome. In addition, in-depth mechanistic knowledge on the intricate and bidirectional interactions between the notochord and the sclerotome are needed to further understand disc morphogenesis.

So far, mechanisms regulating the balance between proliferation and death during IVD development remain largely unknown^43,60^. Previous studies in mouse have shown the importance of SHH signaling but also of the integrity of the notochordal sheath for NCs survival and NP growth^9,29^. Noteworthy, *Jun* gene has been reported to be involved in the regulation of NCs survival at the time of NP formation. Notochord-specific conditional knockout mutant embryos exhibited increased apoptosis in the notochord and not in the sclerotome, reduced cellularity of the IVD, and subsequent fusion of the vertebrae resulting in severe malformations of the axial skeleton^45^. Hippo pathway activity during spine development also warrants further investigation in mice^61^. Genetic analyses have shown that *Tead1* and *Tead2* transcription factor and co-activator YAP protein are essential for maintaining the notochord at E8.5, potentially through the regulation of proliferation and apoptosis^46^. Furthermore, the linked timing of apoptosis and vacuolation in vertebrates’ notochords may reflect apoptosis induction through mechanical stress response to the apparition of vacuoles as discussed in the “turgor pressure-sheath strength” model^19^. This later mechanism provides the notochord with the necessary rigidity for its role as skeletal element for lower-vertebrate embryos development^62^. Previous studies in Zebrafish and Xenopus have shown a correlation between a disorganized extracellular sheath, an impairment in vacuolation and the regulation of apoptosis during notochord development and axis elongation^63–65^. More in-depth functional evaluations are required to elucidate the role of proapoptotic and antiapoptotic members of Bcl-2 family in the notochord along developmental time in order to demonstrate that inhibition of apoptosis would delay or prevent the transformation of the notochord into NP. Further insights into the regulation in NC survival will also help our understanding of the origin and the cause of rare malignant tumors called chordoma^66^. These have been proposed to develop in the axial skeleton from rare intraosseous remnant and dormant notochord cells with prominent vacuoles^67^.

Molecular and genetics studies should be conducted to investigate changes in cell behavior using *in vivo* approaches to assess the impact of genetic alterations in animal models on distinct phenotypic traits in spine development. Although research efforts have improved our understanding of the mechanisms that control NC proliferation in IVD at postnatal stages^68,69^, further studies are required during IVD development to determine cell-autonomous and non-cell-autonomous effects. In addition, recent advanced 3D *in vitro* pluripotent stem cell-based systems modelling axial development in mammals^70,71^ will allow to dissect the effects of signaling pathway activity perturbation, particularly in relation to cellular defects affecting notochord and disc formation. Both *in vivo* and *in vitro* approaches will enhance our understanding of the molecular pathways regulating NC survival, which is also essential to gain deeper insights into both disease mechanisms and regenerative medicine.

### Experimental Procedures

#### Mice

Experiments on mice were conducted according to the French and 138 European regulations on care and protection of laboratory animals (EC Directive 86/609, French Law 2001-486 issued on June 6, 2001). Embryos and Newborns (4 days postnatal, P4) for histology and immunohistochemistry were collected from spontaneous pregnant mice and new-borns respectively from CD1-Swiss background.

#### Histological staining and immunohistochemistry

Embryos from embryonic day E9.5 to E17.5 and post-natal P4 Newborns were collected and were fixed in 4% paraformaldehyde (PFA, Sigma, P6148) at 4°C for 24h and embedded in paraffin (VWR, 10048502) for sectioning in the coronal plane. Spine samples of E17.5 and postnatal P4 were decalcified (EDTA, Euromedex, EU0007, 0.5M pH=7.5 at 4°C) for 7 days before paraffin embedding. Five µm sections were stained following standard protocol with Haematoxylin Eosin Saffron (HES) and Periodic Acid Schiff/Alcian Blue (PAS-AB).

For immunohistochemistry detections, after antigen retrieval, sections were permeabilized (with Triton X-100, Merck T8787, 0.5 % for 20 min) and then the endogenous peroxidase were saturated for 20 min in H2O2 3% (Merck, H1009) in PBS 1X (Vwr, 733-1645). The sections were incubated with the primary antibodies overnight at 4°C after incubation with blocking solution. The next day, the sections were incubated with secondary antibodies conjugated with biotin. Revelation was performed after 45min incubation in streptavidin-peroxidase solution and of DAB substrate (3,3’-Diaminobenzidine). Nuclei were stained with Mayer Haematoxylin (Diapath, C0305), mounted with Eukitt (Kindler GmbH; B0914) and scanned using a Hamamatsu Nanozoomer HT (Hamamatsu Photonics KK) digital scanner at 40X magnification. Table S1 summarizes the dilution and specific buffers for retrieval for each antibody.

#### Immunofluorescence and TUNEL assay

Following collection embryos and mice spines were directly embedded in Super Cryo-embedding Medium (SCEM) (Section Lab) and frozen in isopentane/dry ice, without decalcification and fixation. A specially prepared adhesive film is fastened to the cut surface of the sample in order to support the section while allowing a slow cut. We adapted the Kawamoto’s Film Method^72^ for the use of a Carbone steel blade. Samples were cut along coronal plane and 7μm sections were mounted on slides (CryoStar NX70, Thermo Fisher Scientific) and post-fixed in 2% PFA (2 min) before permeabilization in 0.2% Triton X-100 (in PBS for 10 min and in a blocking solution (10% Foetal Calf Serum, Sigma, G9023; 4% BSA, Sigma, A9647; 0.1% Triton X-100 in PBS) for 1 h. Next, sections were incubated overnight at 4°C with primary antibodies followed by 1 h incubation in secondary antibody (Table S1). Nuclei were stained by Hoechst 33258 pentahydrate (bis-benzimide) (2ug/mL, Thermofisher; H3569) and sections were mounted with ProLong^TM^ (Thermofisher, P36934). Frozen sections for TUNEL staining were post-fixed in 4% PFA for 5 min and cellular membranes were permeabilized with Proteinase K (10μg/mL, Merck P6556) for 5 min. Apoptotic cells were detected using the DeadEndTM Fluorometric TUNEL system (Promega, G3250). Confocal immunofluorescence images were acquired with LFOV FLIM Nikon confocal microscope.

#### Quantification of LAMP1 normalized positive area and cellular density

Quantifications regarding LAMP1 positive area and cellular density were performed on LAMP1 H-DAB images within the open source software QuPath^73^. Images with excessively high or weak DAB signal intensity, as well as NP that were partially out of focus, were excluded from analysis. All measurements were confined to Notochord or NP regions of interest (ROIs), generated using the extension “Segment Anything Model (SAM)”^74^, and manually corrected when necessary. For stages E13.5 and E14.5, note that when the forming NP were connected together with remnant NCs at the pVB level, with no visible separation, they were counted as a single ROI (See Supplementary Fig. S1H). At these stages, regions where the notochord remained rod-shaped were excluded from the quantification. Color deconvolution was adjusted manually and applied to all images, to separate DAB from hematoxylin staining. Nuclei were automatically segmented on the original RGB image, using Stardist^75^ and the “he_heavy_augment” model, with prediction threshold set at 0.5. Detections with median hematoxylin intensity <0.1 or >1.0 were excluded from nuclei count. LAMP1-positive area in μm^2^ was calculated by applying a pixel threshold on the DAB channel. Exported QuPath annotations (ROI and DAB-positive area) and detection counts (nuclei) were analyzed using R (version 4.4.3). For each ROI, cell density was calculated as the total number of detected nuclei divided by the measured ROI area. For normalization, LAMP1 positive area was divided by the number of detected nuclei within the ROI.

#### Quantification and distribution of Caspase-3 signal for apoptosis

Following apoptotic cells detection via immunohistochemistry using Caspase-3 staining, high-resolution images were acquired to visualize notochordal segments along the rostro-caudal axis. Despite careful sample orientation during paraffin embedding, embryonic histological sections often contain interrupted fragments of notochord when the cut is slightly off the strict sagittal plane. Specific regions containing fragment of notochord (NTC) or sclerotome of prospective vertebrae (pVB) or prospective annulus fibrosus (pAF) were manually delineated (See Supplementary Fig. S1A). The quantification strategy allowed the measurement of the number of apoptotic cells among notochord or sclerotome across various areas of interest, enabling distinctions between pVB and pAF. The schematic shown in Fig. S1A summarize the distinct areas counted and classified as i) “Anterior”: including up to two pVBs positioned rostrally relative to the anterior extremity of imaged the notochord fragment, ii) “Posterior”: including up to two pVBs positioned posteriorly relative to the posterior extremity of the imaged notochord fragment and iii) “Bordering the notochord”: defining the pVB and pAF units flanking both extremities of the imaged notochord fragment. Images were analyzed using ImageJ software for region segmentation. For clarity and consistency in spatial classification, colored lines indicated each region of interest on schematic overlays (See Supplementary Fig. S1A). Caspase-3 positive cells were quantified within each region using the CellCounter plugin of ImageJ. A consistent classification was applied across the rostro-caudal axis (for cervical, thoracic, lumbar, caudal regions, Fig. S1B). For reproducibility, quantification was repeated across multiple embryo sections and notochord fragments along the rostro-caudal axis (see Tables 2-3). Data were compiled and statistical analysis performed to assess spatial distribution of apoptotic cells in these defined embryonic domains.

#### Quantification and distribution of Ki-67 signal for proliferation

Following proliferative cells detection via immunohistochemistry using Ki-67 staining, high-resolution images were acquired to visualize notochordal fragments along the rostro-caudal axis. Images were analyzed using ImageJ software for region segmentation. Specific regions containing fragments of notochord seen on the histological section were manually drawn with a dotted line (See Supplementary Fig. S1G). For clarity and consistency in spatial classification, areas of notochord were delineated and colored distinctively according to their location: in blue notochord located at the pVB level and in orange notochord located at the pAF level = future disc area). Ki-67 positive cells were quantified within each areas using the CellCounter plugin of ImageJ. A consistent classification was applied across the rostro-caudal axis (Fig. S1B). For reproducibility, quantification was repeated across multiple E13.5 and E14.5 embryos sections and notochord fragments along the rostro-caudal axis (see Tables 4-5). Data were compiled and statistical analysis performed to assess spatial distribution of proliferative cells in these defined embryonic domains.

#### Statistical analysis

Statistical analyses were performed using R (version 4.4.3) and GraphPad Prism (version 8.0.1). For the distribution of Caspase-3^+^ and Ki-67^+^ cells, multiple groups were compared, a Kruskal-Wallis test was conducted, followed by Dunn’s post hoc test (Dunn, 1964) for pairwise comparisons. To control for the experiment-wise error rate, p-values were adjusted using the Holm method. Pairwise comparisons of normalized LAMP1-positive area were assessed using the Wilcoxon rank-sum test. To correct for multiple comparisons, p-values were adjusted using the Benjamini–Hochberg procedure to control the false discovery rate (FDR). To evaluate the relationship between developmental stage and nuclei density, Kendall’s rank correlation test was calculated using individual ROI. Kendall’s tau coefficient assesses the strength and direction of the monotonic association, with values >0.8 indicating strong, 0.4–0.8 moderate, and <0.4 weak correlation.

---

**Supplementary Fig. S1: Pattern of cell death and schematics for quantification and distribution of apoptosis and proliferation along the rostro-caudal Axis.**

**(A)** Schematic of apoptosis quantification strategy at E12.5, showing defined regions: prospective vertebrae (pVB), annulus fibrosus (pAF), and notochord (NTC). Manually drawn areas (image J) were classified as “anterior,” “bordering the notochord,” or “posterior” based on their position relative to the NTC fragment for the quantification of apoptotic cells and total nuclei. Representation of the counted nuclei for each area using Cell Counter plugin. Black dotted line delimits the classification “Bordering the notochord”. As example, the two red arrows point at pAF unit n°8 and pAF unit n°10 with in between pVB unit n°9 classified in “bordering the notochord” category. **(B)** Whole embryo Caspase-3 staining at E12.5 showing cervical to caudal levels. Scale = 1 mm. **(C-D)** TUNEL staining at E12.5 and at E13.5 with rod-like notochord at the caudal level. Hoechst counterstaining; dotted line outlines the NTC. Scale = 25 µm. **(E-F)** PAS-Alcian Blue and HES stainings at E12.5 showing apoptotic (red arrowheads), vacuolated cells (black arrowheads) and intercellular gap (asterisk) in the NTC.

**(G)** Schematic of proliferation quantification strategy at E12.5. The dotted line indicates the delimitation of the notochord fragment and the colored area the cells counted depending on the region they are located (in blue the pVB and in orange the pAF). **(H)** Schematic of ROI delimitation strategy, for cell density and LAMP1 positive area quantifications. Examples at E13.5 (H) and E14.5 (H’). Yellow line delimitates ROI which includes one (H) or two (H’) NP depending on remnant NC at the level of the pVB (arrows). Scalebars = 50μm. In all panels, sections are rostro-caudal oriented from left-to-right, except panel B (top-to-bottom). NTC: notochord, pAF: prospective annulus fibrosus, pVB: prospective vertebrae.

**Supplementary Fig. S2: Disc development model of the entire rostro-caudal axis**

Spatio-temporal schematic representation of our current working model of IVD development across developmental stages. The top panel shows an overview of IVD morphogenesis and its rostro-caudal progression. Regional differences are particularly noticeable between E13.5 and E14.5, with the cervical and thoracic regions showing more advanced morphological features compared to lumbar and caudal. Changes in cell death, proliferation, and NC phenotype are detailed for the cervical, thoracic / lumbar and caudal regions. At E11.5–E12.5, the notochord is rod-shaped; by E13.5–E14.5, it adopts a periodic swellings or bulges morphology resembling a “string of beads”, becoming elliptical by E17.5 onward. Apoptosis is higher caudally in the pAF sclerotomes while it is uniform along the midline of the notochord at E11.5 and E12.5, forming a central space (dashed line) facilitating ECM deposition. Vacuolated NCs (vNCs) emerge at E11.5. Cytoplasmic vacuole number and size and ECM accumulation increase over time, simultaneously with a general decrease in cell density and NP volume expansion. Arrows indicate proposed physical cues from pAF apoptosis (straight) and pressure constraint from forming pVB (curved), both guiding NC expansion into pAF, along with the increase of proliferative cells in this region. ECM: extracellular matrix; pAF and pVB: prospective annulus fibrosus and vertebrae. Created in https://BioRender.com.

**Supplementary Table S1: Immunohistochemistry and immunofluorescence detailed procedures**

## Acknowledgments

The authors would like to thank A. Henry for the initiation of the project and N. Wyckens for technical advice. We acknowledge M. Ambroset from the MicroPICell core facility (SFR Bonamy, BioCore, Inserm UMS 016, CNRS UAR 3556, Nantes, France), member of the Scientific Interest Group (GIS) Biogenouest, IBISA, and the national infrastructure France-Bioimaging supported by the French national research agency (ANR-24-INBS-0005 FBI BIOGEN), the SC3M facility from SFR Francois Bonamy, University of Nantes and the UTE-IRS-UN Animal facilities staff for animal care, breeding, and for their investment in the project.

